# Untangling the cell immune response dynamic for severe and critical cases of SARS-CoV-2 infection

**DOI:** 10.1101/2021.03.23.436686

**Authors:** Rodolfo Blanco-Rodríguez, Xin Du, Esteban Hernández-Vargas

## Abstract

COVID-19 is a global pandemic leading high death tolls worldwide day by day. Clinical evidence suggests that COVID-19 patients can be classified as non-severe, severe and critical cases. In particular, studies have highlighted the relationship between the lymphopenia and the severity of the illness, where CD8^+^ T cells have the lowest levels in critical cases. In this work, we aim to elucidate the key parameters that define the course of the disease deviating from severe to critical case. To this end, several mathematical models are proposed to represent the dynamic of the immune response in patients with SARS-CoV-2 infection. The best model had a good fit to reported experimental data, and in accordance with values found in the literature. Our results suggest that a rapid proliferation of CD8^+^ T cells is decisive in the severity of the disease.

## 1. Introduction

COVID-19 caused by SARS-CoV-2 infection is a global pandemic which has caused more than 40 millions confirmed cases and more than 1 million deaths worldwide. People of all age can be infected where around 20% of the cases are asymptomatic, 60% appear with mild or moderate conditions, and 20% are severe or critical cases [2]. Most of the countries have taken emergency actions, these actions include confinement of their population, travel restrictions, forced use of mask in public spaces, and even a nighttime curfew. In this situations, epidemiological models have been key to mitigating COVID-19 pandemic and many others [15].

There are three coronaviruses (SARS-CoV, MERS-CoV and SARS-CoV-2) that can cause pneumonia, which can be fatal. SARS-CoV-2 is transmitted mainly via respiratory droplets, the median incubation period is around 4 days before symptom onset [13], most of symptomatic patients developing symptoms within 11.5 days [21]. The viral load reaches its peak within 5-6 days of symptom onset [32]. An animal model using rheus macaques reported two peaks of viral RNA, the first peak is input of the virus, while the second one is due to authentic viral replication [49].

Most of COVID-19 patients present without any symptoms or only mild respiratory symptoms [5]. Moderate cases present principally fever, cough, and fatigue; less common symptoms are sputum production, headache, hemoptysis, and diarrhea [18]. Most of moderate patients are recovered, however, a portion of these patients are hospitalized. Approximately 20% of cases develop severe illness, requiring intensive care unit (ICU) treatment because of complications, including acute respiratory distress syndrome, arrhytmia, and shock. Critical patients have symptoms of dyspnea and they are more likely to be older [44]. The bulk of patient who die had comorbidities like hypertension, heart disease, dibetes, among others. Respiratory failure is the most common cause of death, followed by sepsis, cardiac failure, hemorrhage, and renal failure. In these cases lymphopenia, neutrophilia and thrombocytopenia were usually observed [48].

A key determinant factor of disease severity in SARS-Cov-2 is age, in particular individuals over 65 years have the greatest risk of requiring intensive care [5]. As other viral infections, the severity of the disease in the elderly is not directly attributed to the viral titter but to the host immune response [17]. Severe patients are characterized by difficult in breathing and low blood oxygen level; in some cases there even be secondary infection by bacteria and fungi that may cause respiratory failure, which is the cause of death in most fatal COVID-19 cases [5, 40].

The storm of cytokines released by immune system in response to the infection can result in sepsis that is the cause of death in 28% of fatal COVID-19 cases [48]. For influenza infection, adaptive immune response against viral infection impairs innate immune defense against bacterial infection [30]. Immune therapies inhibiting viral infection and regulation of dysfunctional immune response are key to block pathologies [40, 9, 37].

It is still controversial whether virus persistence can increase the severity of the disease. SARS-Cov-2 viral dynamics has shown remarked differences between severe patients and non-severe patients. Viral load peak is higher in non-severe patients (~ 10^8^ copies/mL) than severe patients (~ 10^7^ copies/mL). Also, viral shedding time has been longer in severe patients [39], even the virus is detectable until death [51]. It also has been reported the mean viral load of severe cases around 60 times higher then mild cases [23].

Similar to other viral infection, adaptive immune response have a key role in SARS-CoV-2 infection, particulary T cells [17]. It remains unclear weather T cell response are helpful or harmful in COVID-19. Mathew et al. [27] identified three immunotypes revealing different patterns of lymphocytes response in hospitalized COVID-19 patients. Immunotype 1 was associated with highly activated CD4^+^ and CD8^+^ T cells; immunotype 2 had less CD4^+^ T cell activation; and immunotype 3 had lacked activated T and B cell response. Mortality ocurred for patients with all three immunotypes. On the other hand, patients with severe conditions have shown lymphopenia associated with COVID-19, where CD8^+^ T cells have a major impact [6, 28]. Direct virus killing lymphocytes could be part of the problem, as SARS-CoV particles have been found in T cells, monocytes and macrophages [12].

There are several studies around the dynamic changes of lymphocytes [25, 22], showing low level of lymphocytes in severe patients. In [47], Zhang et al. analyze the dynamic changes of lymphocyte subsets and specific antibodies in coronavirus disease. They obtained blood samples of 707 patients from Wuhan, China, which were classified into moderate, severe and critical groups. Their results shows that the counts of total T cells, CD4^+^ T cells and CD8^+^ T cells were significantly decreased with the increased severity of illness. The levels of these lymphocytes could be helpful markers to indicate the severe illness of COVID-19 and to understand the pathogenesis of COVID-19.

Mathematical modeling can be pivotal to dissect the dynamics between severe and non-severe COVID-19 patients. Epidemiological models have been of great help to follow the pandemic evolution, evaluate the results of different scenarios and reveal the health measures that could help to mitigate the pandemic [1, 36, 24]. Similarly, mathematical models at within-host level can help us to understand viral infections and the immune response [33, 16, 45, 11, 8].

While these in-host models [33, 16, 45] are fitted to viral data from COVID-19 patients to infer the interaction with the immune response, there has not been any study to quantify the differences between severe and critical patients with COVID-19. As far as we know, there are no models fitted to T cells data in order to examine the relation between T cell dynamic and severity of the illness. In this work, we contribute to the mathematical study of SARS-CoV-2 dynamic and the T cell dynamics to elucidate the principal role of lymphocytes in the develop of the disease between severe ad critical patients.

## 2. Materials and methods

### 2.1. Experimental data details

Here we considered the data reported by Zhang et al. [47]. They collected from 707 COVID-19 patients in Wuhan, China between February and April, 2020. The patients were classified into moderate, severe and critical groups. The moderate cases were those with fever, typical symptoms and pneumonia. 206 severe cases had respiratory distress, blood oxygen saturation less than 93%, or arterial partial pressure of O_2_ to fraction of inspired oxygen ratio less than 300 mmHg. 91 critical cases had respiratory failure, shock or multiple organ dysfunction needing intensive care unit treatment.

The counts of total T cells, CD4^+^ T cells, CD8^+^ T cells, B cells, and Natural Killer (NK) cells were analyzed with FACSCanto flow cytometer from 50 *μ*1 of whole blood. The patients were 48.5% males and 51.5% females. Most of the patients had fever, cough, expectoration, shortness of breath, chest distress, diarrhea at the illness onset. The most common comorbidities of the cases were hypertension (37.9%), diabetes (17.1%) and cardiovascular disease (11.1%). There were 30 deceased. The total T cells, CD4^+^ T cells and CD8^+^ T cells in moderate patients were relatively stable compared to those for several and critical cases. The severe and critical group had a lower count of lymphocyte from the illness onset but gradually recovered to the normal levels. More details can be found in the original paper [47]. The data are displayed in Fig. 1, the median is represented as points and the dashed lines represent the interquartile range (IQR). Data are reproduced from the original paper [48] using plotDigitizer.

**Figure 1:**
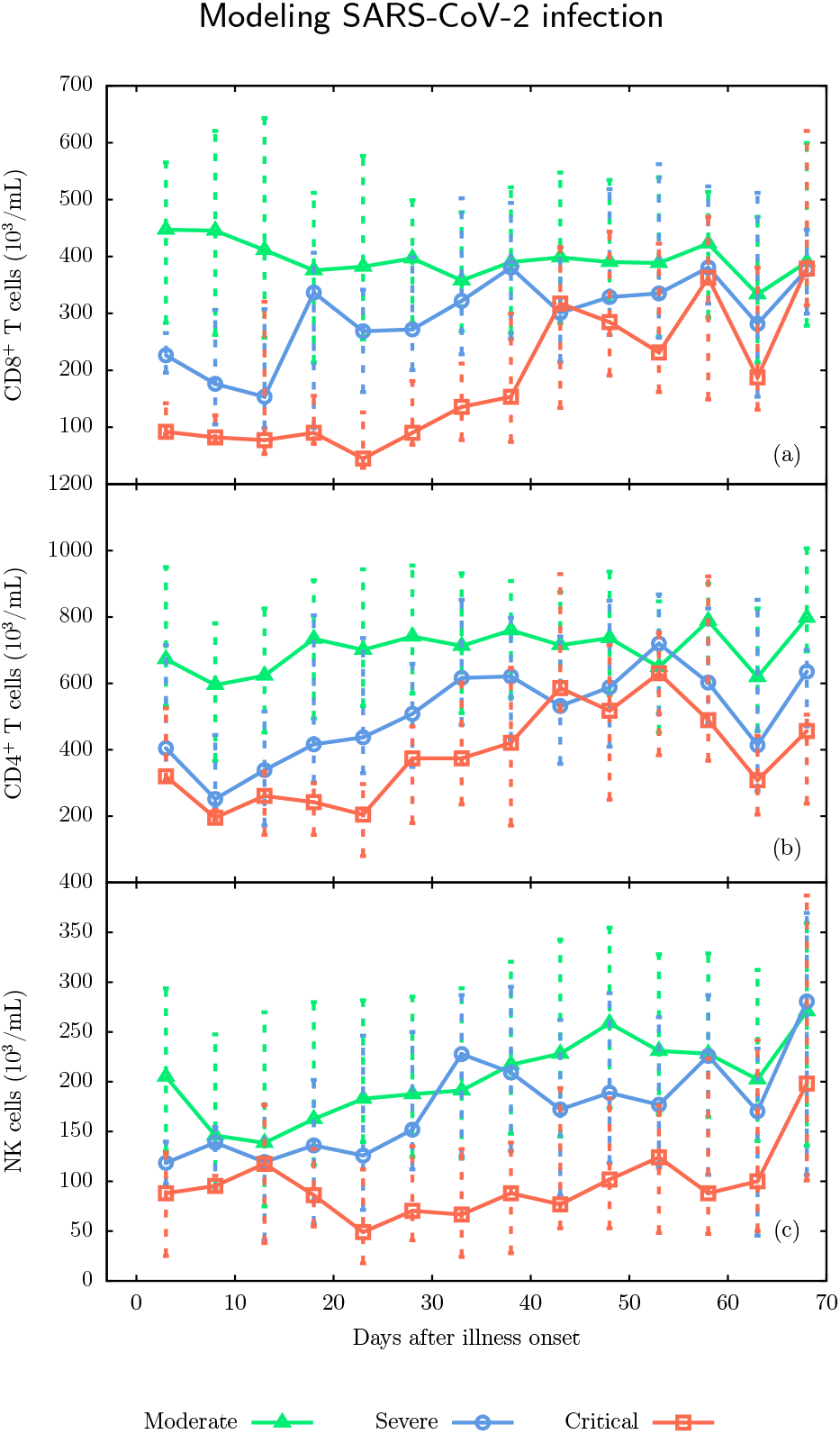
Experimental data. (a) CD8^+^ T cells, (b) CD4^+^ T cells and (c) natural killers cells levels. Data are reproduced from reference [47].

### 2.2. Mathematical model

In [16] has been reported a mathematical model to represent the interaction between SARS-CoV-2 infection and immune response dynamics. The model is given by:

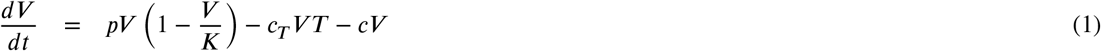

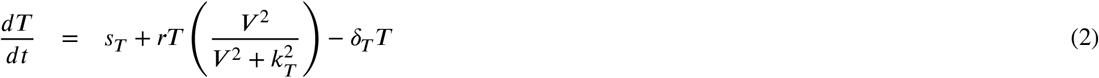

where *V* is the virus level, *T* the number of CD8^+^ T cells, *p* the viral replication rate with maximum carrying capacity *K*, and *c* the rate of cleared virus. *c_T_VT* represent the rate of killing of infected cell by the immune response. In this model is assumed that the activation of T cell proliferation by *V*, at a rate *r*, follows a log-sigmoidal form with half saturation constant *k_T_*. The parameter *s_T_* = *δ_T_T*(0) represent T cell homoeostasis with *δ_T_* as the half life of T cells and *T*(0) the initial number of them. In figure 2 is shown a schematic representation of this model.

**Figure 2:**
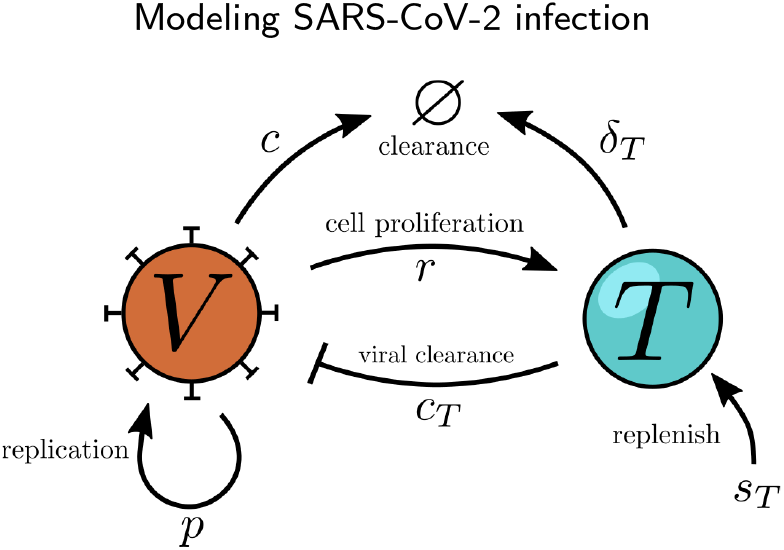
Schematic representation of the viral infection model with immune response. Virus (*V*) induces CD8^+^ T cells (*T*) a rate *r* which inhibits the viral replication through the clearance of the infected cell a rate *c_T_*. *T* cells are replenished with constant rate *s_T_* and die with a rate *δ_T_*. Virus are replicated with a rate *p*. The rate of clearance of the virus due to processes not directly related to immune system is represented by *c*.

Contrary to [16], in this work we fitted the data of CD8^+^ T cells to the model (2) using the median of the CD8^+^ T cells count for severe and critical cases from [47]. We reduced the number of parameters to be identified to *p*, *c_T_*, and *r*. The half-life of CD8^+^ T cells in humans have been estimated from 34 days [29] to 255 days [43]; therefore, we take *δ_T_* = 0.01 day^-1^. We fix *c* = 2.4, *K* = 10^8^, and *k_T_* = 1.26 × 10^5^ for both cases; this values were taken in accordance to [16]. Also we use the initial viral level *V*(0) = 0.31 copies/ml. The *s_T_* parameter for each case was fixed with the respective initial value *T*(0). Due to lack of data before illness onset, we assumed the initial level of T cells equal to the median of the CD8^+^ T cells in the day 3 (*T*(0) = *T*(3)) from the reported data for each case. Infection time was assumed at −3 days after illness onset (daio).

In our model, we included CD8^+^ T cells in the peripheral blood of patients described above, who have a wide range of comorbidities. We only considered several and critical cases, since data from moderate cases showed no marked changes during the disease course.

Similar to CD8^+^ T cells, there is a evidence of activation and/or exhaustion markers at CD4^+^ T cells [6]. Even it has been suggested that CD4/CD8 ratio is significantly higher in critical patients than non-critical patients [31]. Because of that, we modified the model. We considered that CD4^+^ helps to proliferation of CD8^+^ T cells which occurs at rate *αT*_4_ where *T*_4_ is CD4^+^ T cell level and *α* is a free parameter to be estimated. We use piecewise linear fits to generate a time-dependent function *T*_4_(*t*) using the experimental data displayed in Fig. 1b. We also explored different ways to integrate CD4^+^ T cell data to our model, however, we do not obtain good results.

Furthermore, natural killer cells (NK) are critical in the first-line defense against viral infection, and integrate innate and adaptive immune responses [42]. It has been correlated the number and function of NK during SARS-CoV-2 infection with the severity of the disease [50, 26]. Therefore, we explored the viral clearence due to NK (*N*) at rate *c_N_VN*. Similar to modification above, we use piecewise linear fits to generate *N*(*t*) using data in Fig 1c.

### 2.3. Parameter estimation

The ordinary differential equations of the model were solved using a Dormand-Prince fifth-order Runge-Kutta algorithm. The estimation of the free parameters was performed by minimize the Root Mean Square Error (RMSE) using the difference between the experimental measurement (*y_i_*) and the predictive output (*ȳ_i_*) as follows:

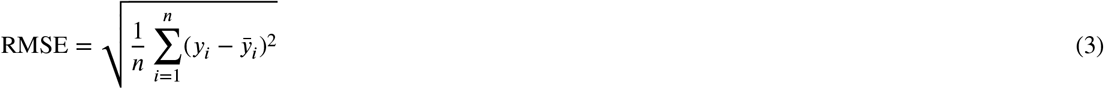

where *i* is the corresponding sample and *n* is the total number of measurement. To minimize the RMSE we used the Differential Evolution (DE) algorithm [38]. We implemented a DE algorithm using GPU parallelization with code written in CUDA-C, this implementation accelerate the optimization ten times at least. Some details about a DE implementation on GPU can be found in [41].

### 2.4. Akaike information criterion

In order to compare between different models, we used the Akaike information criterion (AIC) defined by:

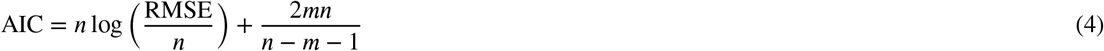

where *n* is the number of data points and *m* is the number of unknown parameters. A lower AIC values means a better description of the data.

### 2.5. Identifiability analysis

A mathematical model is said to be identifiable when the parameter set can be uniquely determined. Here we used the profile likelihood method proposed by [35]. In this method one by one the parameters are set to a range of values centered at the optimized value; the other parameters are re-optimized using the same cost function, which is the RMSE above mentioned. This methodology can detect both structurally and practically non-identifiable parameters [14]. Structural non-identifiability is related to the model structure and practical non-identifiability takes into account the amount and quality of the data. A parameter can be identified when the profile likelihood presents a concave form. However, if the data are insufficient and manifest large variability, the parameter could be practically non-identifiable. This can be visualized as a relatively flat valley in the profile likelihood. A structurally non-identifiable parameter has a profile that maintains a constant RMSE when the parameter is varied.

### 2.6. Bootstrap

The experimental data displayed in Fig 1(a) present a highly variable response to SARS-CoV-2 infection, hence we performed bootstrap fits to mimic a stochastic environment of the infection. Notice that the quartiles in Fig 1(a) are asymmetric, that is, that the median values of the experimental data are not in the middle of the IQR. For that reason, we generated 27 discrete data between lower limit and upper limit of the IQR (including the median value) for each point of the daio (x-axis). We then performed a nonparametric bootstrap approach using Monte Carlo resampling. Data were resampled with replacement to have a sample of equal size to the generated values above. The parameters are estimated from the resampling. We adapted our DE code to perform 100 parameter estimations at the same time using GPU parallelization in order to save computational time. A total of 1000 optimizations (10 runs of our DE code) were performed using different sets of resampled data. We obtained the corresponding parameter distribution from refit our model in each of these repetitions.

## 3. Results

The experimental data for CD8^+^ T cells and their respective fit of the model above are displayed in Fig. 3a. The experimental data were reproduced from [47]. For severe cases, the CD8+ T cell response starts about 10 to 20 daio reaching its peak between 35 to 45 daio, while for critical the CD8+ T cell response starts late, around 30-40 daio with a peak between 40 to 50 daio. Note that critical cases begin with a lower level of CD8^+^ T cells than severe cases (half of them), however, both reaches approximately the same level of the moderate cases at the end of the disease course. The total count of cells (accumulative sum) between both cases are in the same order of magnitude although is lower for critical cases (1.3 × 10^7^) than severe cases (2.2 × 10^7^).

**Figure 3:**
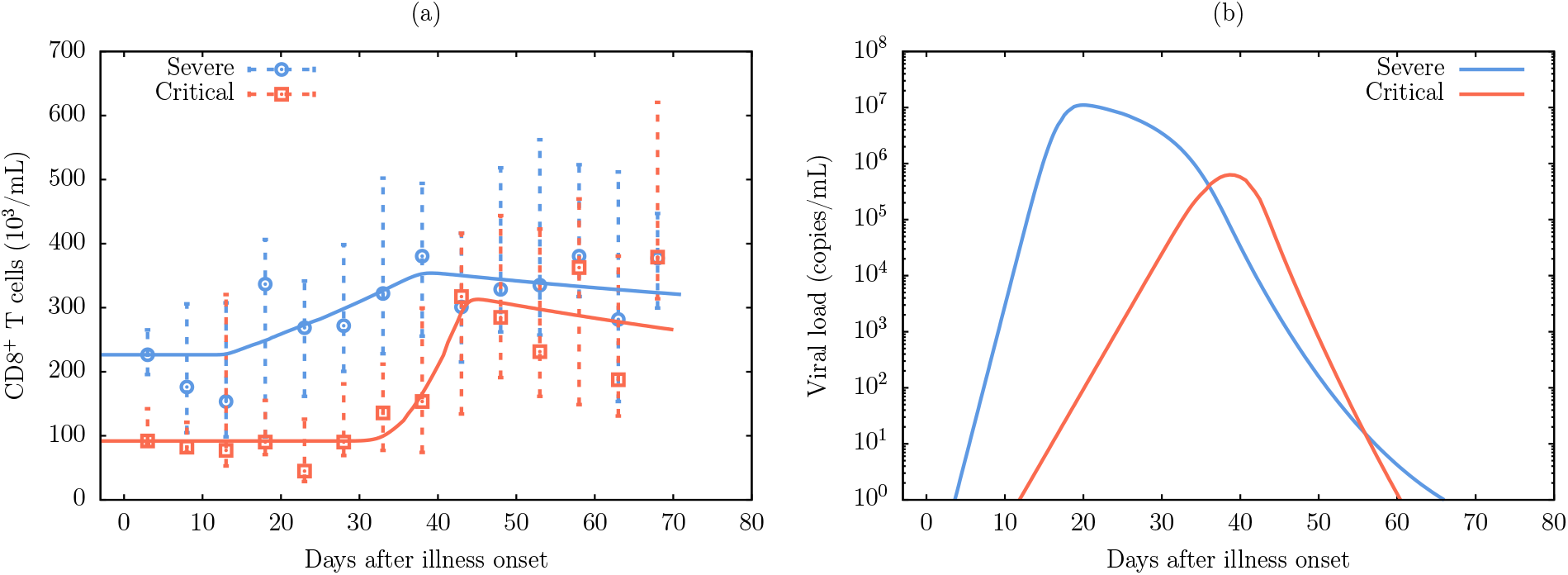
(a) Model fitted to experimental data for severe and critical cases. The continuous line are simulation. Points are the median of the experimental data and dashed lines are the interquartile range. (b) Viral load obtained from the model after the parameter optimization for severe and critical cases. Data are reproduced from reference [47].

The viral load obtained using the model with the parameter fitted to CD8^+^ T cells experimental data is displayed in Fig. 3b. The viral load peaks around 40 daio for critical cases and 20 daio for severe cases. There is a delay in the peak of the viral load for critical cases compared to severe cases; critical viral load peak is two orders of magnitude lower than severe viral load peak. This result is consistent with those reported in [39], although in those results the difference is of one order of magnitude between severe and non-severe. The total viral count is higher for severe case (1.2 × 10^8^) than critical cases (4.5 × 10^6^).

The profile likelihood analysis was performed with the unknown model parameter. These profiles are shown in Figs. 4a, 4b and 4c. Critical cases show a minimum for viral replication rate *p* and viral clearance *c_T_* implying identifiability of these parameter. The likelihood profile for the CD8^+^ T cell proliferation rate *r* does not show a minimum for critical cases. In severe cases, the profiles for *c_T_* and *p* show a minimum while for *p* it is not very clear. This results suggest the possibility to infer model parameters from experimental data reported.

**Figure 4:**
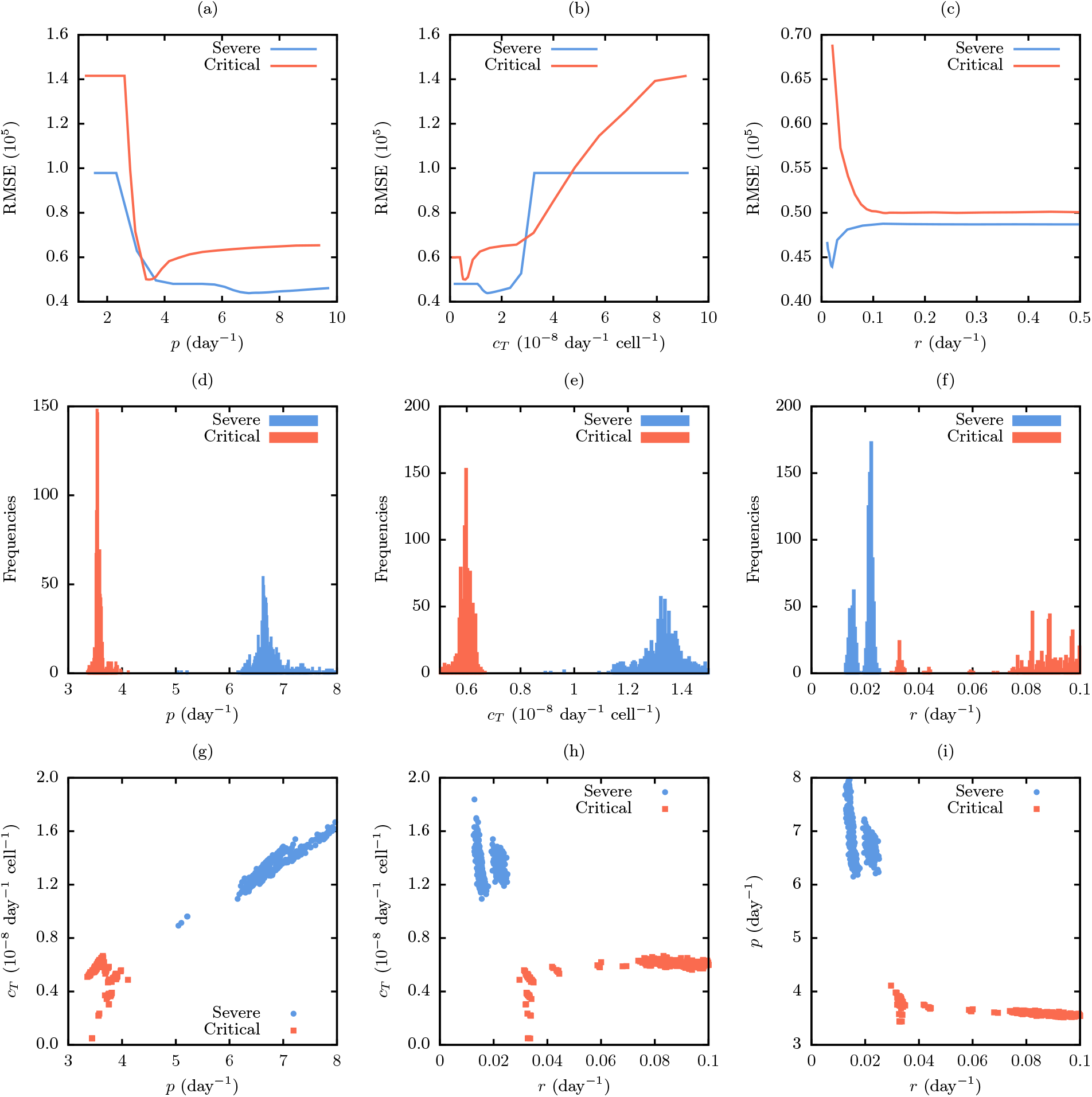
Profile likelihood for the model parameters; (a) viral replication rate *p*, (b) viral clearance *c_T_* and (c) CD8^+^ T cell proliferation rate. Parameter distribution from 1000 bootstrap fits: (d) *p*, (e) *c_T_* and (f) *r*. Parameter ensembles from bootstrap: (g) *p-c_T_*, (h) *r-c_T_* and (i) *r*-*p*.

The best fitted parameters are presented in Table 1 for CD8^+^ T cells and the two cases of illness severity. The viral replication rate *p* for critical cases is a half of that for severe cases. The viral clearance *c_T_* for critical cases is approximately one third of that for severe cases. The CD8^+^ T cell proliferation rate *r* is higher for critical cases. These results suggest that the rapid proliferation of T cells and the low clearance rate are key in the development of the disease.

**Table 1.** Model parameter values using CD8^+^ T cells experimental data from [47]. Fixed parameters are taken from [16].

| Fixed parameters |  |  |  |
| --- | --- | --- | --- |
| $K$ | $10^8$ copies/mL | | |
| $c$ | $2.4 \text{ day}^{-1}$ | | |
| $k_T$ | $1.26 \times 10^5$ copies/mL | | |
| $\delta_T$ | $0.01 \text{ day}^{-1}$ | | |
| Critical cases |  |  |  |
| Parameter | Best fit | Median | CI (95%) |
| $p \text{ [day}^{-1}\text{]}$ | 3.50 | 3.55 | (3.45 - 3.81) |
| $c_T \text{ [} 10^{-8} \text{ day}^{-1} \text{ cell}^{-1}\text{]}$ | 0.596 | 0.595 | (0.384 - 0.636) |
| $r \text{ [day}^{-1}\text{]}$ | 0.131 | 0.097 | (0.033 - 0.137) |
| Severe cases |  |  |  |
| Parameter | Best fit | Median | CI (95%) |
| $p \text{ [day}^{-1}\text{]}$ | 6.99 | 6.67 | (6.29 - 7.63) |
| $c_T \text{ [} 10^{-8} \text{ day}^{-1} \text{ cell}^{-1}\text{]}$ | 1.47 | 1.34 | (1.16 - 1.54) |
| $r \text{ [day}^{-1}\text{]}$ | 0.020 | 0.021 | (0.014 - 0.024) |

Due to high variability of the data, we performed bootstrap fits in order to obtained the confidence interval of the model parameter estimated. Figs. 4d, 4e and 4f shows the distribution in parameter values for severe and critical cases. The three free parameter show clear difference between cases. The viral replication rate *p* in critical cases decreased 45% with respect to severe cases, the rate of killing of infected cell by immune response *c_T_* decreased 53% for critical cases; whereas that CD8^+^ T cell proliferation rate *r* was four times the rate for severe cases. Table 1 shows the median and 95% confidence interval of each parameter. The median values presented for critical cases are consistent with the values for the best fit, except for *r*. Severe cases median values shows the opposite behavior, only *r* is in accordance with the value for the best fit. This discrepancy could due to high variability of the experimental data used.

In order to explore the dependencies of the parameters, we displayed scatter plots in Figs 4g, 4h and 4i. These plots reveal that there are no strong correlation between *p*, *c_T_* and *r*. However, we can notice a slight inter-dependence between *p* and *r* parameters for critical cases; and *p* and *c_T_* parameters for severe cases. In the former, increasing *r* decrease *p*; and in the latter, increasing *p* increase *c_T_*.

We explored the modification of the model by adding CD4^+^ T cells and NK responses. Fitting these models to the data revealed that including CD4^+^ T cells as a helper of the proliferation of CD8^+^ T cells does not improve the fits, similarly for NK response. Increasing the number of parameters to optimize does not improve the results neither. In the Table 2 is shown the AIC number for each model. The parameters *α* and *c_N_* tend to small values (10^-3^ and 10^-10^ respectively), leading the same dynamics as the first model. In the third and fourth model we added CD4^+^ T cell helper as a log-sigmoidal form, however, in these models the viral load does not reach to be cleared; notice that they have the highest AIC numbers.

**Table 2.** Model comparison

| Model | Fit | Severe |  | Critical |  |
| --- | --- | --- | --- | --- | --- |
|  |  | RMSE | AIC | RMSE | AIC |
| $\dot{V} = pV(1 - V/K) - c_T VT - cV; \dot{T} = s_T + rT(V^2/(V^2 + k_T^2)) - \delta_T T$ | $p, c_T, r$ | 4.38 | 57.33 | 4.98 | 58.12 |
| $\dot{V} = pV(1 - V/K) - c_T VT - cV; \dot{T} = s_T + rT(V^2/(V^2 + k_T^2)) - \delta_T T + \alpha T_4$ | $p, c_T, r, \alpha$ | 4.34 | 61.32 | 4.94 | 62.11 |
| $\dot{V} = pV(1 - V/K) - c_T VT - cV; \dot{T} = s_T + rT(T_4^2/(T_4^2 + k_4^2)) - \delta_T T$ | $p, c_T, r, k_4$ | 7.94 | 64.99 | 6.40 | 63.68 |
| $\dot{V} = pV(1 - V/K) - c_T VT - cV; \dot{T} = s_T + rT(V^2/(V^2 + k_T^2))(T_4^2/(T_4^2 + k_4^2)) - \delta_T T$ | $p, c_T, r, k_4$ | 8.05 | 65.07 | 13.7 | 68.29 |
| $\dot{V} = pV(1 - V/K) - c_T VT - cV - c_N VN; \dot{T} = s_T + rT(V^2/(V^2 + k_T^2)) - \delta_T T$ | $p, c_T, r, c_N$ | 4.28 | 61.24 | 5.00 | 62.19 |

## 4. Discussion

The role of the immune system during SARS-CoV-2 infection is fragmented. The T cell kinetics seem to be decisive in the resolution of severe or non-severe patients [22]. CD8^+^ T cells are relevant for killing infected cells during viral infections [34]. Furthermore, a defective immune response may lead to further accumulation of immune cells in the lungs causing overproduction of cytokines, resulting in a cytokine storm leading to multi-organ damage [40, 10]. Therefore, an abnormal proliferation of T cells could lead to critical state to the COVID-19 patient. Quantification of the dynamics of these T cells could help to identified the critical cases in the early stage of the disease.

A recent study [3], patients with hematologic cancer show that higher CD8+ T cell counts is associated with improved survival. Also, robust CD4+ T cell response in conjunction with a diminished CD8+ T cells is key in survival patients. Our simulations highlight a clear difference between the parameters that model critical case and those that model severe cases. The principal difference is in the rate of T cell proliferation *r*, this rate is high in critical cases. This is in accordance with [20, 46], suggesting a hyperactivation and overaggressive CD8^+^ T cell response. However, it is still unclear whether the T cells in COVID-19 patients are exhausted or just highly activated [6].

On the other hand, fitting results show a viral clearance rate *c_T_* for severe cases is higher than that for critical cases. The viral replication rate *p* for severe cases is also higher than that for critical cases which translates to a higher viral peak. Therefore, although the severe cases have a low production of CD8^+^ T cells compared with that critical cases, these cells clear a major viral load. Notice that viral shedding takes approximately the same time for both types of cases, this suggests that critical cases may have a dysfunctional immune response where there are excessive infiltration of T cells which could cause widespread inflammation and multi-organ damage; infected cells are slowly cleared.

In [47], both critical and severe cases begin do not have any significant difference for CD4+ T cell levels. Nevertheless, there is a tendency of low levels of CD4+ T cells in critical patients. Note that in normal conditions, IL-2 improve transcription of FOXP3 which is widely recognized as a suppressor of the T cell response. However, in severe COVID-19 patients, activated T cells fail to express FOXP3 [19]. This T cell dysregulation promotes prolonged inflammation and tissue destruction. Model selection was not able to show that CD4+ T cells play an important role in viral clearance or CD8+ T cell proliferation.

Here, we explored the innate immune response against viral infection adding to our model NK cell response, nevertheless, this did not improve the AIC with just CD8^+^ T cell response. This may be attributted that NK cells play an important role in the begining of the infection, such dynamics can not be capture in the used data set [47]. It has been reported that the upregulation of human inhibitory receptors is one more strategy that SARS-CoV-2 uses to modulate NK cell cytotoxicity. It is clear that increase expression of such receptors lead to NK cell exhaustion and decrease their ability to clear viral infection. The mechanisms that drive NK cell exhaustion are poorly understood [4]. Several upregulated genes in peripheral blood from COVID-19 patients are involved in the apoptosis pathways, suggesting lymphopenia is due to apoptosis by SARS-CoV-2. In SARS-CoV-2 infection NK cells exit the peripheral blood, contributing to local inflammation and injury. NK cells that remain in the circulation show an exhausted phenotype that facilitate virus spread [26].

The model that best describe the data is considering CD8^+^ T cell response. The best fits show a delay viral peak for critical cases, the difference is 20 days; while CD8^+^ T cells levels peak approximately in the same time for both cases and with almost the same level. Critical cases have the T cell response peak 5 days after their viral load peak. Finally, it is worth to mentioning that experimental data taken in [47] have samples with different comorbidities and ages, this results could be different if we took data from homogeneous group.

Figure 3(a) does show that CD8+ T cells and the virus start to grow earlier in the severe patients respect to the critical one. However, while the CD8+ T cell is delayed in the critical patients, it reaches similar levels than the severe patients. This difference between severe and critical COVID-19 patients can be attributed to the effects of aging to the immune systems which is highly altered during viral infections [17]. Previous mathematical modeling work [17] had formulated that the slower viral growth presented in aged individuals may lead to less immune stimulation [7].

